# Adaptive dynamics of sexual signaling and discrimination under different mating conditions

**DOI:** 10.1101/2025.07.23.666195

**Authors:** Benjamin Allen, Samantha E. Rothberg, Brian A. Lerch

## Abstract

A wide variety of animals engage in same-sex sexual behavior (SSB). Some instances of SSB appear to arise from individuals mating indiscriminately without regard to the sex of their partners. Theory suggests that indiscriminate mating strategies can be selected if sexes are not completely distinguishable—that is, without strong signals of sexual identity. We model the coevolution of sexual signaling and discrimination using the adaptive dynamics approach to long-term trait evolution. Our model relaxes two major assumptions from past phenomenological theory. First, we consider a mechanistic mating process with tunable encounter rates and mating costs. Second, we vary reproductive investment roles to allow for better correspondence with empirical examples of SSB. We confirm that coevolution of sexual signaling and discrimination can lead to two distinct equilibria: one with no sexual signals and indiscriminate mating; the other with perfect signaling and exclusively different-sex sexual behavior. Selection for indiscriminate mating is strongest at very low or very high encounter rates. Mating costs lead to selection for reduced overall mating rate, but this can lead to either more or less SSB, depending on how this reduction is achieved. Overall, our model highlights the importance of ecologically relevant parameters for the evolution of SSB.

## Introduction

Same-sex sexual behavior (SSB) occurs in a wide range of animals, including echinoderms, mollusks, insects, birds, reptiles, and mammals (Bagemihl, 1999; Bailey and Zuk, 2009; Roughgarden, 2004; Scharf and Martin, 2013; Sommer and Vasey, 2006). SSB takes a variety of forms, including mounting, courtship, and pair bonding. The prevalence of SSB has likely been underreported, since many instances are presumably overlooked, misidentified, or never published (Bagemihl, 1999; Bailey and Zuk, 2009; Rice et al., 2013; Russell et al., 2012).

Since SSB does not directly produce offspring, biologists have sought to explain its evolution, leading to many non-mutually exclusive hypotheses. Some hypotheses include potential adaptive benefits of SSB, such as social bonding (de Waal, 1995; Mann, 2006) or intrasexual competition (Kureck et al., 2011; Lane et al., 2016; Preston-Mafham, 2006). Others propose that SSB arises as a byproduct of pleiotropy (Han and Brooks, 2015; Logue et al., 2009) or from a lack of discrimination with regard to mating partners (Bailey and French, 2012; Hoving et al., 2012; Sales et al., 2018).

Incomplete sex discrimination—or “broad mating filters” (Richardson and Zuk, 2023)—may be understood as a response to the fitness costs of missing a potential mating when sexes are not completely distinguishable (Hoving et al., 2012; Lerch and Servedio, 2021). The extreme case of indiscriminate sexual behavior (ISB) has recently received much interest (Lerch and Servedio, 2021, 2023), especially in light of recent work suggesting that early sexually reproducing animals likely mated indiscriminately (Monk et al., 2019).

Less attention has been paid to the possibility that the distinguishability of the sexes may itself evolve. Evidence that the traits of potential mating partners influence the expression of SSB (Han et al., 2016; Mizumoto et al., 2022; Richardson and Zuk, 2023) makes clear that sex recognition requires a detectable signal or cue. Signals of sexual identity—visual, auditory, olfactory, behavioral, or otherwise—are themselves subject to evolution, both influencing and influenced by the evolution of sex discrimination.

A recent theoretical model (Lerch and Servedio, 2023) analyzed the coevolution of sexual signaling and sex discrimination. In this model, signaling and discrimination increase efficiency in mating effort, but are subject to fitness costs that are treated phenomenologically as free parameters. They find that signaling and discrimination coevolve in a positive feedback loop. If both traits are present initially in sufficient frequency, they reinforce one another, leading to strong sexual signals and strong sex discrimination. On the other hand, they find that the absence of both sexual signals and sex discrimination is always a potential selective equilibrium. This bistability of mating strategies may have implications for understanding both ancestral and contemporary sexual behaviors among clades of animals. In particular, it suggests that transitions from ancestral ISB to exclusively different-sex behavior (DSB) are not inevitable, but arise only under particular conditions.

It remains unclear, however, how the coevolution of signaling and discrimination depends on key parameters of the mating process. First, mating costs (mortality, energetic, or other material costs of mating or courtship) have been proposed as an important variable in the evolution of SSB (Stojkovic et al., 2010) and estimated empirically (Fedorka et al., 2004; Martin and Hosken, 2004; Pereira et al., 2010; Stojkovic et al., 2010), but not incorporated into previous models. Second, population density and frequency of encounters are argued to influence the expression of SSB, but baseline expectations for the direction of this influence are lacking. Hoving et al. (2012) argue that infrequent encounters should lead to selection for weaker sex discrimination, because the opportunity costs of missed matings are very high under these conditions. In contrast, Weldon (2005) and Scharf and Martin (2013) suggest that high density conditions (frequent encounters) may select for highly responsive males that mate quickly and therefore discriminate less effectively. Theory is needed to tease apart these arguments. Third, asymmetric investment in reproduction may play a role—empirically, the clearest examples of imperfect sex discrimination (Hoving et al., 2012; Richardson and Zuk, 2023; Sales et al., 2018) are in males, which are the lower-investing sex in these species. Elucidating the effects of these variables requires a more mechanistic representation of the mating process.

Here, we develop a new coevolutionary model of sex discrimination and sexual signals. Our model explicitly represents the key biological variables of encounter rate, mating and signaling costs, sex roles, and sex ratio as model parameters. There are two sexes, “Signalers”: who may produce a signal to inform others of their sex, and “Finders” who choose whether or not to initiate mating with others they encounter. The signal quality of Signalers, and the level of sex discrimination of Finders (i.e. breadth of the mating filter; Richardson and Zuk, 2023), are continuously evolvable traits. Using the adaptive dynamics approach (Dercole and Rinaldi, 2008; Dieckmann and Law, 1996; Metz et al., 1996), we analyze the long-term coevolution of signal quality and discrimination strength over many rounds of mutant invasion and fixation. We use this model to assess the effects of key ecological parameters on the selective landscape of indiscriminate and discriminate sexual behavior.

## Model

### Signaling and discrimination

We consider a large, sexually reproducing population of size 2*N*. A fraction *s* of the population are Signalers, while the remaining fraction, 1 − *s*, are Finders. Signalers may, or may not, produce a signal to inform others of their sex. Finders seek out mates, possibly taking signals into account when deciding whether to initiate mating. Fertilization and reproduction occur only from mating between a Signaler and a Finder. Only Finders initiate mating; consequently, SSB can only occur between Finders.

There are two evolvable traits: the signal quality, *q*, of Signalers, and the discrimination strength, *d*, of Finders. A Signaler’s value *q* equals the probability that a Finder can detect this Signaler’s sex. Thus, a Signaler with *q* = 1 is always identified as a Signaler, while a Signaler with *q* = 0 is indistinguishable from Finders. A Finder’s value of *d* is the probability that a Finder will *not* initiate mating in the absence of a signal. Thus, a Finder with *d* = 0 mates indiscriminately with all others it encounters, while one with *d* = 1 mates exclusively with those it detects to be Signalers. Aggregating over the probabilities of detection and mating (Fig. **1**), the overall probability that a given Finder with discrimination strength *d* initiates mating is

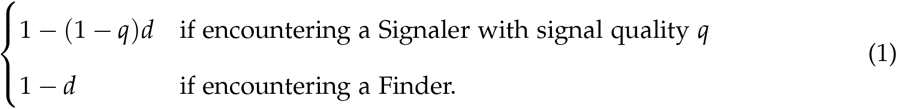

**Figure 1:**
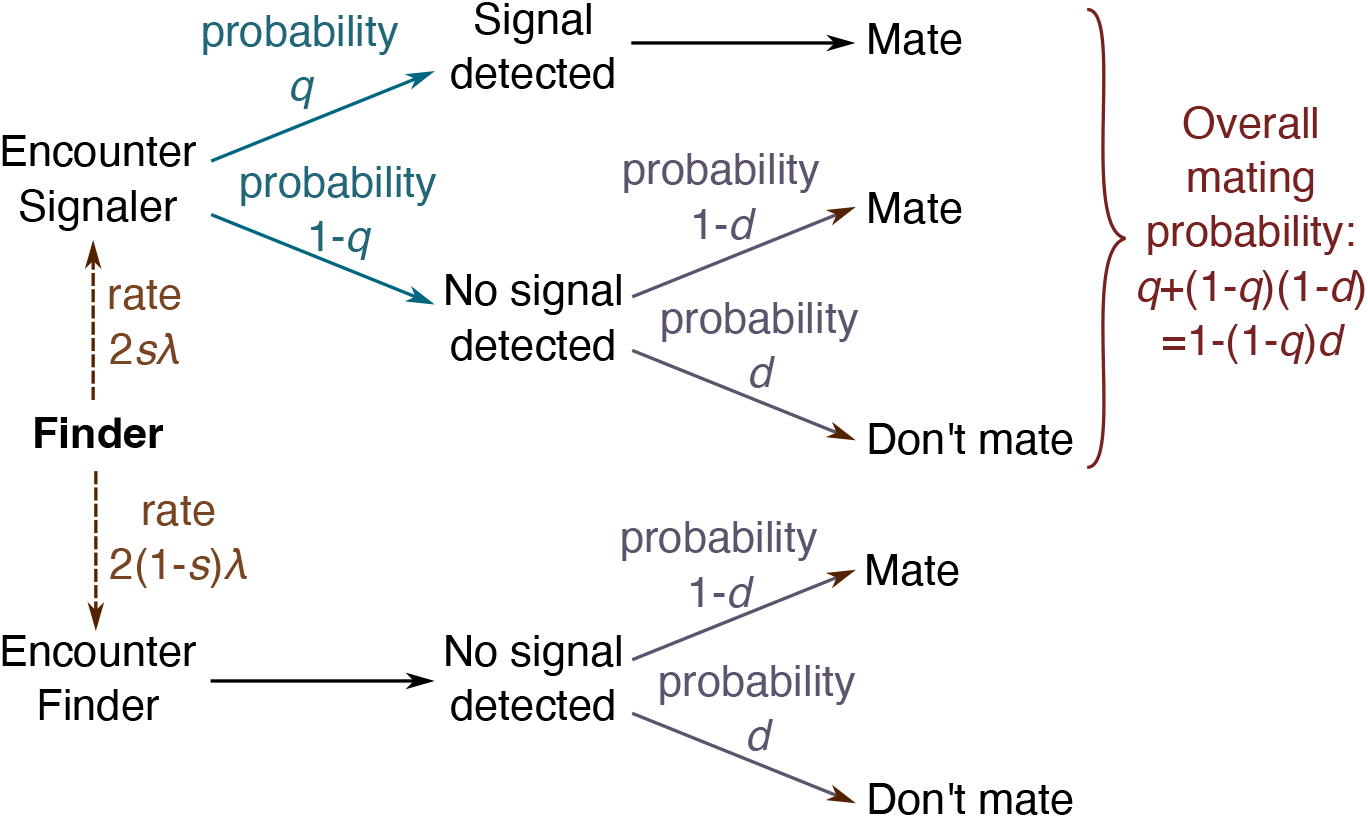
Probability tree for sexual signaling and discrimination. A Finder encounters each other individual at rate *λ*/*N* per season. Since the are 2*N* individuals, of which a fraction *s* are Signalers, each Finder encounters Singalers at rate 2*sλ* and Finders at rate 2(1 − *s*)*λ*. When encountering a Signaler, the Finder detects a signal with probability equal to the signal quality *q*. If a signal is detected, the Finder always initiates mating; otherwise, the Finder initiates mating with probability 1 − *d*, where *d* is the Finder’s discrimination strength. The overall probability of mating in this encounter is therefore *q* + (1 − *q*)(1 − *d*), which simplifies to 1 − (1 − *q*)*d*. When encountering another Finder, no signal is detected, and the Finder initiates mating with probability 1 − *d*.

It is worth clarifying our use of the term “signal”, which can be used with different implications in sexual selection theory. In our model, signaling has a binary outcome (detected or not), and carries no information other than the sex of the Signaler. The signal does not, for example, reveal information about Signalers’ other traits; nor does it help Finders to locate Signalers. Furthermore, whether a Signaler’s sex is detected depends only on their signal quality, and not on any traits of Finders.

### Mating and reproduction

Time is divided into seasons, each representing a window of opportunity for mating. Within each season, time is continuous. Encounters between individuals are modeled as Poisson processes. Specifically, each Finder encounters each other individual at rate *λ*/*N* per season, where the parameter *λ* tunes the encounter rate. For each encounter, the Finder initiates mating with probability according to Eq. (1). Once initiated, mating cannot be declined by either party.

We initially assume that Signalers are high-investment, meaning they can be fertilized at most once per season, while Finders are low-investment and may fertilize any number of Signalers. A Signaler is fertilized the first time they mate with a Finder in a given season, and cannot be fertilized again in the same season. However, a fertilized Signaler remains available to Finders for (nonreproductive) mating. A Finder, in contrast, fertilizes as many Signalers as they are the first to mate with in a season. At the end of each season, each fertilized Signaler reproduces, resulting in a fitness gain of *b* to each parent (i.e., 2*b* surviving adult offspring, on expectation and weighted by reproductive value if necessary).

Later, we will consider the role-reversed case of high-investment Finders and low-investment Signalers. In most—but not all—cases, the low-investment sex corresponds to males and the high-investment sex to females (Janicke et al., 2016). Correspondingly, our baseline case typically applies to male-male SSB, and the role-reversed case to female-female SSB.

### Costs

Mating and signaling incur costs, which are measured in units of reproductive fitness. Each mating (regardless of whether it results in fertilization) comes at cost *c*_F_ to Finders and *c*_S_ to Signalers. We require that *b* > *c*_F_ and *b* > *c*_S_, so that reproduction results in a net fitness gain to each parent.

Additionally, each Signaler pays a cost *C*(*q*) to produce their signal of quality, *q*. We assume that the signaling cost *C*(*q*) is nondecreasing in *q* (that is, *C*′(*q*) ≥ 0 for all *q*), and that *C*(0) = 0 (no signal incurs no costs).

In addition to the above costs, the population size is kept constant by a density-dependent background mortality rate that is equal across all individuals.

### Invasion fitness and adaptive dynamics

We model long-term coevolution of sexual signaling and discrimination using the adaptive dynamics approach (Dieckmann and Law, 1996; Metz et al., 1996). We assume that mutation is rare, so that the population is typically monomorphic with regard to *q* and *d*. Occasional mutants arise, with signal quality, *q*′, or discrimination strength, *d*′, differing from the resident values of *q* and *d*. The success of such a mutant is determined by invasion fitness functions (Dieckmann and Law, 1996; Metz et al., 1996), which quantify the growth rate of a rare mutant trait value (i.e. *q*′ or *d*′) in the context of resident values *q* and *d*. We derive these invasion fitness functions here in the case of even sex ratio (*s* = 1/2). Uneven sex ratios are considered in the Appendix.

Consider a rare mutant Signaler type, whose signal quality, *q*′, differs from the resident signal quality, *q*. Such a Signaler encounters Finders at total rate *λ* per season, and each encounter leads to mating with probability 1 − (1 − *q*′)*d* according to Eq. (1), so that this Signaler mates *λ* 1 − (1 − *q*′)*d* times per season, on expectation. According to the Poisson process, the probability that this mutant Signaler is fertilized (i.e. mates at least once) in a season is 1 − *e*^−*λ*(1 − (1 − *q′*)*d*)^. The resulting in expected reproductive benefit *b (*1 − *e*^−*λ*(1 − (1 − *q′*)*d*^) is offset by costs *c*_S_*λ*(1 − (1 − *q*′)*d*) from mating and *C*(*q*′) from signaling. Aggregating benefits and costs, the invasion fitness of the mutant Signaler is

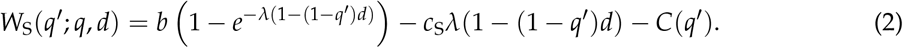

It is not necessary to include a term for background mortality, since this is equal across types and therefore does not affect relative fitness.

Next, consider a rare mutant Finder type whose discrimination strength, *d*′, differs from the resident discrimination strength, *d*. To successfully reproduce, a mutant Finder of this type must be the first to mate with a given Signaler in a given time unit. A given Signaler experiences mating attempts at rate (*λ*/*N*)(1 − (1 − *q*)*d*′) from each mutant Finder, and rate (*λ*/*N*)(1 − (1 − *q*)*d*) from each resident Finder. The probability that the mutant Finder mates with a given Signaler in a given season, prior to any other mating with this Signaler, is calculated as

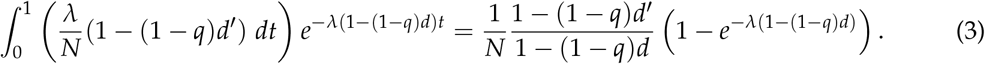

This is multiplied by the number, *N*, of Signalers, and the benefit, *b*, of each fertilization, to obtain the expected reproductive benefit to the mutant Finder. Meanwhile, each mutant Finder incurs mating costs of *c*_F_*λ*(1 − (1 − *q*)*d*′) from mating with Signalers, and *c*_F_*λ*(1 − *d*′) from mating with Finders. Aggregating, the invasion fitness of the mutant Finder type is

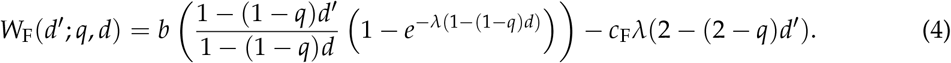

Under the additional assumption that mutations are incremental (i.e., of small effect), the coevolution of signal quality, *q*, and discrimination strength, *d*, is modeled using the canonical equation of adaptive dynamics (Dieckmann and Law, 1996):

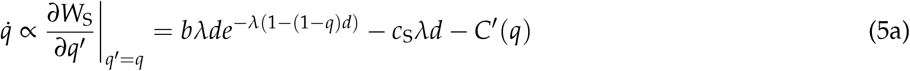

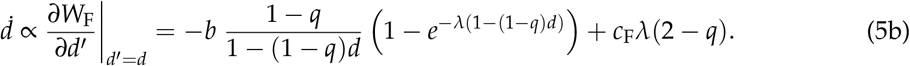

Above, the symbol ∝ denotes “proportional to”. The constants of proportionality in Eqs.(5a) and (5b) are *Nµ*_S_*σ*_S_ and *Nµ*_F_*σ*_F_, respectively, where *µ*_S_ and *µ*_F_ are mutation rates, and *σ*_S_ and *σ*_F_ are expected mutational step sizes, in the respective traits, *q* and *d*. These constants affect evolutionary trajectories and rates of evolution, but not the trait values or stability of evolutionary equilibria (see below).

## Results

We find there are two possible equilibria to which traits may converge. One, with *d* = *q* = 0, represents purely indiscriminate sex behavior (ISB) and no signaling. The other, with *d* = 1 and *q* close to or equal to one, represents purely different-sex sexual behavior (DSB) and high-quality signaling.

Specifically, the pure ISB state, *d* = *q* = 0, is a stable equilibrium if *C*′(0) > 0 (signaling is costly even at low quality) and *b*(1 − *e*^− *λ*^)/*λ* > 2*c*_F_ (see Appendix B.1). The latter condition ensures that, for Finders, the expected benefits of mating exceed the costs, taking into account that, with even sex ratio and no discrimination, half of matings are with the same sex. Under these conditions, ISB is stable because there is no advantage to creating a signal—which would be ignored—nor to discriminating on the basis of a nonexistent signal.

The pure DSB state, *d* = *q* = 1 is a stable equilibrium if (*be*^− *λ*^ − *c*_S_)*λ* > *C*′(1) (Appendix B.2). This condition weighs the net costs and benefits to Signalers of maintaining perfect signal quality, under full discrimination. If this condition is met, DSB is stable because reducing signal quality would result in missed mating opportunities, while reducing discrimination strength would incur unnecessary mating costs. If not, a stable DSB equilibrium with *d* = 1 can still arise, with an intermediate equilibrium signal quality *q* <1 that balances the need to mate at least once per season with the costs of excess (nonreproductive) mating for Signalers.

In cases for which both equilibria are stable, the evolutionary outcome depends on the initial population state. Each stable equilibrium has a basin of attraction, with all trajectories initiating in each basin ultimately converging to the corresponding equilibrium. The curve separating these basins of attraction contains an unstable saddle point (Appendix B.3). Bistability between ISB and DSB equilibria was also found by Lerch and Servedio (2023) under a different model.

By analyzing how these basins of attraction expand or shrink with changes in parameters, we shed light on how key biological variables such as population density, mating costs, and sex ratio affect selection for discriminate or indiscriminate sexual behavior. Our findings are summarized in Table 1.

**Table 1:** Effect of parameters on basins of attraction.

| Parameter name | Symbol | Increase favors |
| --- | --- | --- |
| Encounter rate | $\lambda$ | First DSB, then ISB* |
| Finders' mating cost | $c_F$ | DSB |
| Signalers' mating cost | $c_S$ | ISB |
| Marginal cost of signal quality | $C'(q)$ | ISB |
| Fraction of Signalers | $s$ | ISB |

\* Effects of $\lambda$ are nonmonotonic in the baseline investment-role case. For the role-reversed case, increase in $\lambda$ favors DSB.

### Effects of encounter rate

The encounter rate, *λ*, is positively related to population density, a key empirical variable that is thought to affect the evolution of sex discrimination. However, there is so far no consensus as to the direction of the effect (Hoving et al., 2012; Scharf and Martin, 2013; Weldon, 2005).

When *λ* is small—meaning encounters are rare—most initial conditions lead to the ISB equilibrium (Fig. **2**A). This is because, when potential mates are scarce, any opposite-sex mating opportunity may be one’s last. In this case (and as long as *b* > 2*c*_F_, which guarantees sufficient reproduction to offset mortality), the danger of missing a mating opportunity outweighs the cost of possible nonreproductive matings. For very small *λ*, Eq. (5a) for 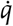 is dominated by the − *C*′(*q*) term, so that signal quality evolves downward and the DSB equilibrium is lost. This supports Hoving et al.’s (2012) “shot in the dark” hypothesis that low population density favors ISB.

As the encounter rate *λ* increases, the basin of attraction of the ISB equilibrium first shrinks, then grows, for the following reasons. First, as encounters become more frequent (Fig. **2**B–C), there is less downside to missing a potential mating opportunity, since other opportunities may arise in the same time-step. Consequently, discriminate strategies become increasingly viable (allowing Finders to avoid excess mating costs), leading more initial conditions toward the DSB equilibrium. However, when encounters become so frequent that Signalers readily attain their maximum of one fertilization per time-step (Fig. **2**D–F), the benefit of high-quality signaling is reduced relative to its cost. This reduced benefit leads to selection for reduced signal quality, which in turn makes discrimination less efficacious, and the ISB basin of attraction widens again, ultimately eliminating the DSB equilibrium.

**Figure 2:**
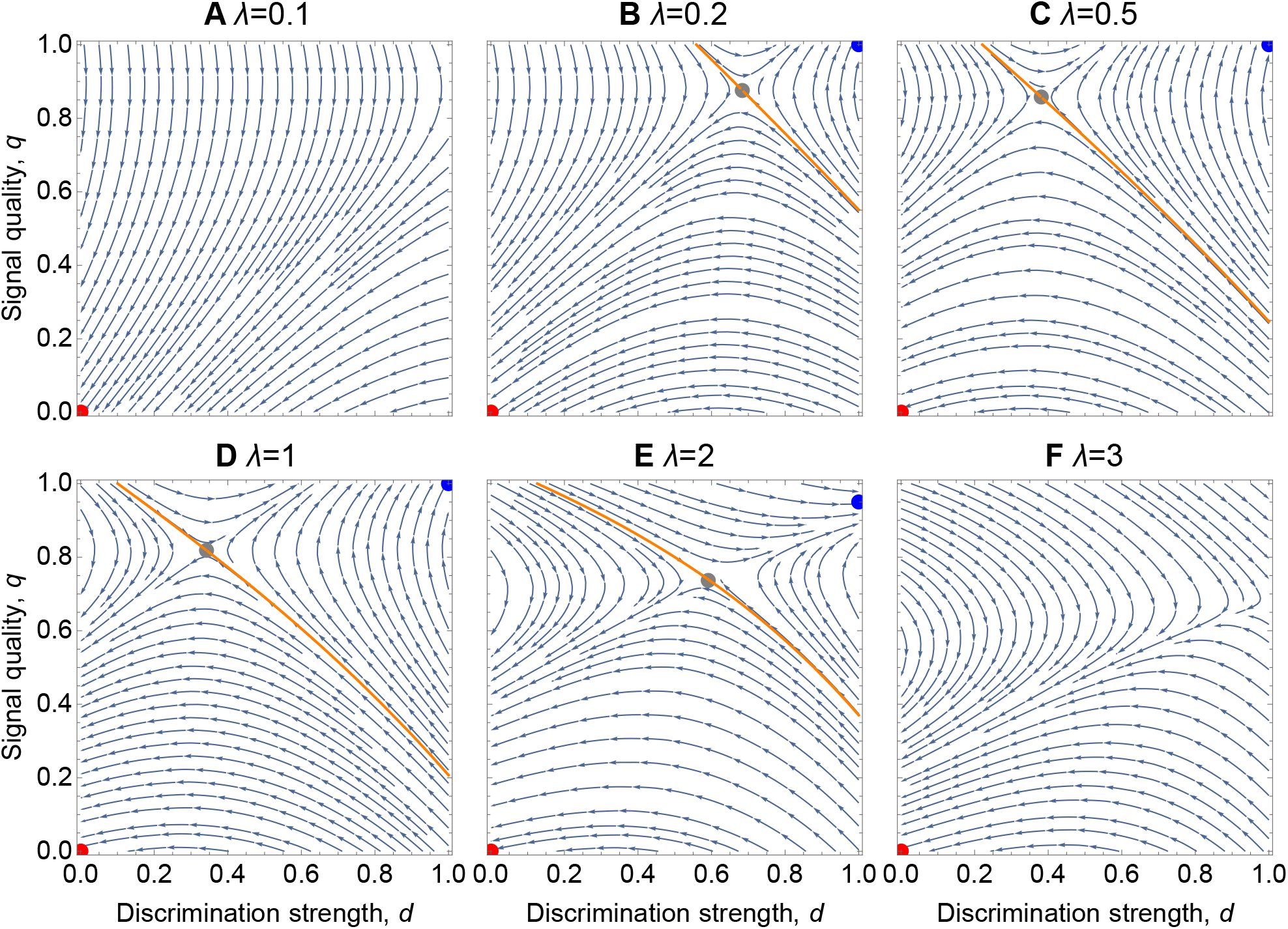
Effect of encounter rate *λ* on coevolutionary dynamics of signaling and discrimination. Stream plots show the adaptive dynamics of signal quality, *q*, and discrimination strength, *d*, based on Eq. (5), with constants of proportionality set to one. Other parameters are fixed at *b* = 1, *c*_F_ = *c*_S_ = *C*′(*q*) = 0.1. Stable ISB (indiscrminate sexual behavior) and DSB (different-sex sexual behavior) equilibria are marked with red and blue points, respectively, while a gray point indicates an unstable interior fixed point. The orange curve shows the separatrix between the basins of attraction of the two stable equilibria, below which the population evolves to the ISB equilibrium and above which the population evolves to the DSB equilibrium. **A** For very low encounter rates, all trajectories lead to ISB equilibrium (*d* = *q* = 0). **B–C** As the encounter rate increases, a DBS fixed point emerges and its basin of attraction expands. **D–F** As the encounter rate increases further, the DSB basin of attraction shrinks until all trajectories lead to ISB again.

### Effects of mating and signaling costs

For Finders, the only benefit of sex discrimination is avoidance of excess mating costs. Consequently, if Finder mating costs are negligible (*c*_F_ = 0; Fig. **3**, top row), there is no selective advantage to discrimination, and most trajectories converge to a stable ISB equilibrium. As Finder mating costs increase (Fig. **3**, top to bottom), avoidance of nonreproductive mating becomes more advantageous. This leads to a stable DSB equilibrium with widening basin of attraction as *c*_F_ increases.

**Figure 3:**
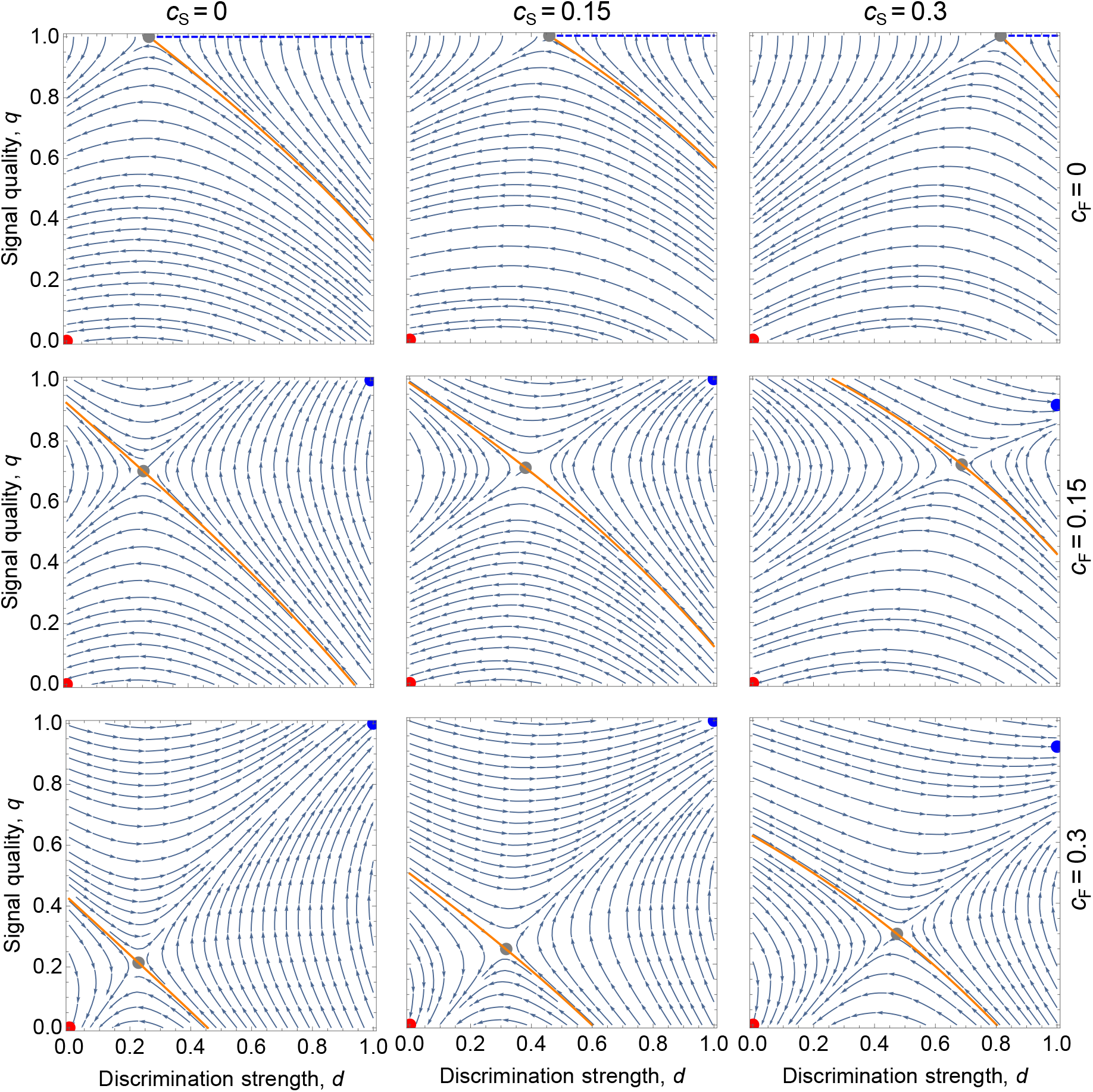
Effects of mating costs. Varying the costs of mating for Finders, *c*_F_, and Signalers, *c*_S_ lead to opposite effects. If mating is costless for Finders (*c*_F_ = 0; top row), there is no selection pressure to discriminate (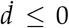 always). This leads to an ISB equilibrium with large basin of attraction, and a line segment of neutrally stable equilibria with perfect signaling. As mating becomes more costly for Finders (top to bottom), selection pressure to avoid nonreproductive mating leads to a stable DSB equilibrium with expanding basin of attraction. In contrast, as mating becomes more costly for Signalers (left to right), selection favors reduced signal quality to avoid excess mating. This makes discrimination less effective, so the ISB basin expands. Other parameters are fixed at *b* = *λ* = 1 and *C*(*q*) = 0.1*q*.

Notably, the Signaler mating cost, *c*_S_, has the opposite effect (Fig. **3**, left to right). As *c*_S_ increases, selection favors avoidance of excess mating via reduction of signal quality (i.e. less distinguishability from Finders). In turn, this downward selection on signal quality reduces the effectiveness of discrimination, causing the ISB basin of attraction to expand.

Overall, for both Finders and Signalers, increased mating costs leads to selection for reduced mating rate. However, this is achieved differently for the two sexes. Finders reduce their mating rate by increasing discrimination strength, which reduces SSB; whereas Signalers reduce mating by decreasing signal quality, which increases SSB.

Unsurprisingly, high signaling costs also counteract the advantages of signaling. The higher the marginal cost *C*′(*q*) of increasing signal quality, the less advantageous it is to send high-quality signals, leading more trajectories toward ISB (Supplementary Fig. **S1**).

### Effects of sex ratio

To probe the effects of uneven sex ratio, we vary the fraction, *s*, of Signalers in the population. Adaptive dynamics equations for arbitrary sex ratio are derived in Appendix A. We find (Supplementary Fig. **S2**) that the ISB basin of attraction expands as the fraction of Signalers increases, and contracts as *s* decreases. This is because, under a Finder-biased sex ratio (*s* <0.5), selection favors discrimination in order to avoid nonreproductive Finder-Finder matings. Conversely, under a Signaler-biased sex ratio (*s* > 0.5), there is less advantage to discrimination, since Finders mostly encounter the opposite sex anyways. Similar patterns regarding sex ratio were found by Lerch and Servedio (2021).

### Effects of investment roles

Above, we have assumed that Signalers are high-investment, and Finders low-investment, so that SSB is confined to the low-investment sex (usually males). Here we explore the role-reversed case of high-investment Finders and low-investment Signalers. This means that a Finder can reproduce at most once per season, whereas a Signaler can reproduce as often as it is the first to mate with a given Finder in a given season. In the role-reversed case, SSB is confined to the high-investment sex (usually females).

Assuming even sex ratio, the canonical equation of adaptive dynamics in the role-reversed case becomes

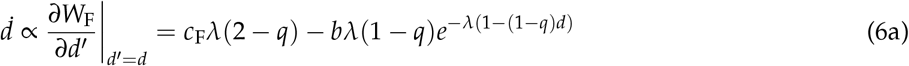

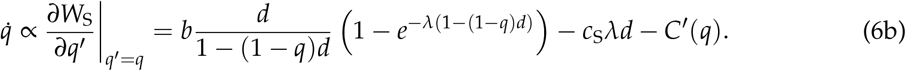

See Appendix A.2 for the derivation and generalization to arbitrary sex ratio.

For low-to-moderate encounter rates (Fig. **4**A–C), dynamics in the role-reversed case resemble those in the baseline case. Indeed, in the limit of small encounter rates (*λ* → 0), the role-reversed case is mathematically equivalent to the original case (see Appendix C). Moreover, in both investment-role cases, increasing *λ* from zero has the initial effect of shrinking the ISB basin of attraction, by reducing the downside of missing a mating opportunity and thereby increasing the viability of discriminate strategies.

**Figure 4:**
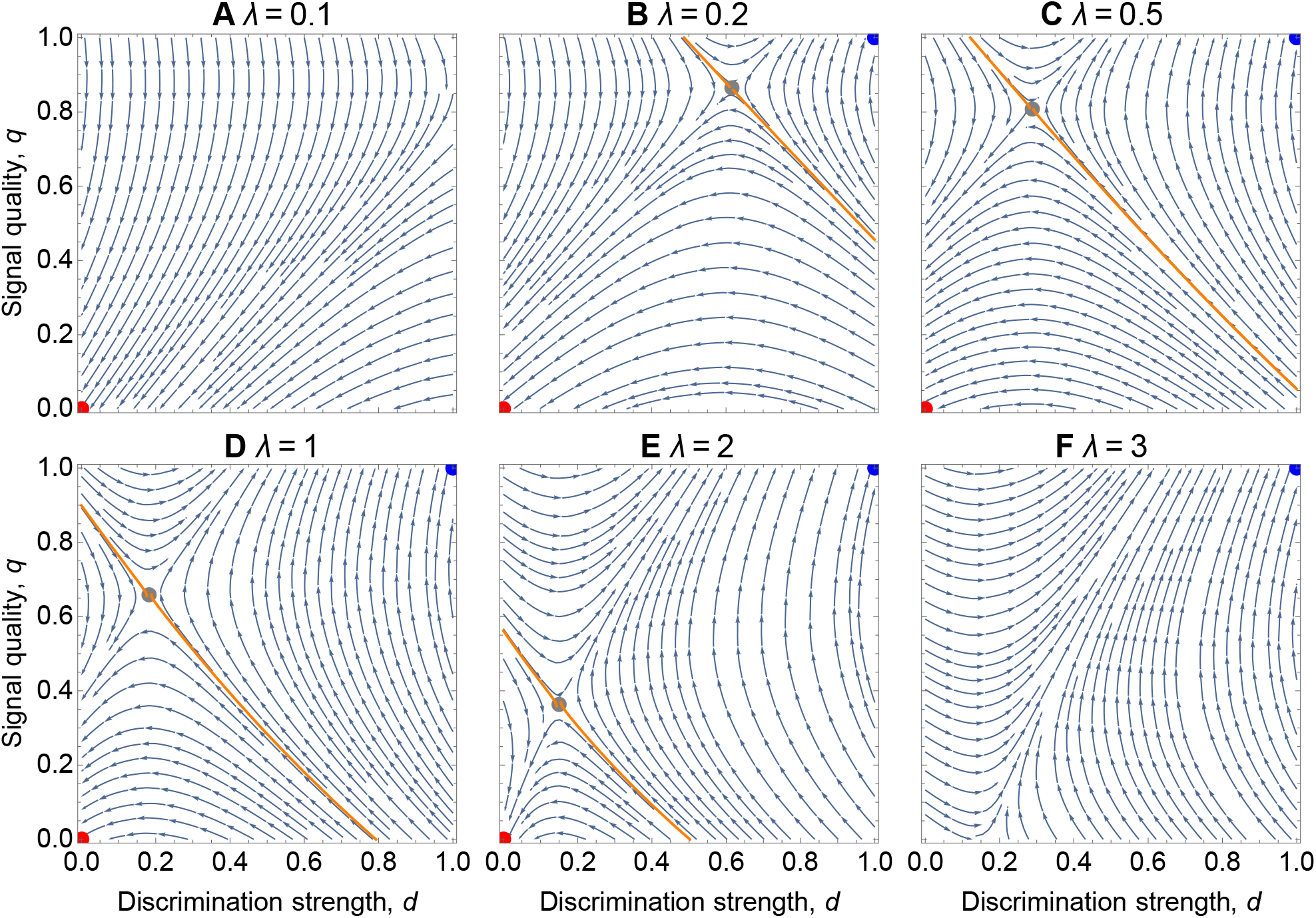
Effects of encounter rate *λ* with investment roles reversed. In this case, Finders are high-investment and Signalers low-investment. **A** As in the baseline case (Fig. 2A), all trajectories lead to ISB when encounters are sufficiently rare. **B.F** However, in contrast to the non-reversed case, the basin of attraction of DSB expands monotonically as the encounter rate increases. This is because (i) the opportunity costs of a missed mating decrease, and (ii) Finders evolve increased discrimination to avoid excess mating (with either sex). Other parameters are fixed at *b* = 1, *c*_*F*_ = *c*_*S*_ = *C*’(*q*) = 0.1.

However, differences arise between the investment-role cases as the encounter rate increases further (Fig. **4**D–F). When *λ* is large enough that Finders easily attain their maximum of one reproductive mating per time-step, selection favors increased discrimination strength to avoid the costs of excess matings (which are nonreproductive beyond the first). Thus, in contrast to the baseline case, increasing encounter rate unidirectionally tilts the dynamics in the role-reversed case toward DSB. As with mating costs, this difference emerges because of the mechanisms by which excess mating is reduced: stronger discrimination for Finders versus reduced signal quality for Signalers.

## Discussion

### Stability of discriminate and indiscriminate sexual behavior

Our work adds to theoretical arguments (Lerch and Servedio, 2021, 2023; Monk et al., 2019) and empirical evidence (Hoving et al., 2012; Richardson et al., 2024; Richardson and Zuk, 2023) that indiscriminate or partly discriminate sexual behavior may be an adaptive response to imperfect information. The quality of information (i.e. signals) may itself coevolve with sex discrimination or the lack thereof. The result is that sex discrimination evolves to balance the costs of nonreproductive mating (which may be small) against the opportunity costs of missed reproduction.

We find that, for a broad range of parameter values, both indiscriminate sexual behavior (ISB) and exclusively different-sex sexual behavior (DSB) are evolutionarily stable. Depending on initial conditions, a population may converge to a state in which sexes are indistinguishable and mating is indiscriminate, or one with clearly identifiable sexes and exclusively opposite-sex mating. Each equilibrium state, once established, is robust to invasion by alternative strategies (see also Lerch and Servedio, 2023). In this light, both discriminate and indiscriminate mating strategies are consistent with theory.

### Evolution of sexual identity signaling

The bistability typically exhibited by our model is driven by positive coevolutionary feedback between signaling and discrimination. When signaling and discrimination are both absent, there is no advantage to either behavior. Conversely, when signaling and discrimination are both perfect, reduction of either trait becomes disadvantageous.

Empirical studies of SSB have typically focused on males, who in these contexts are the party that initiates mating (the Finders, in our model). Our work makes clear the importance of considering how members of the other sex signal their sexual identity. Finders must base their mating decision off of some sexually dimorphic traits, but the traits that individuals use to differentiate the sexes are rarely considered in the context of SSB (Lerch and Servedio, 2023). The few empirical studies which have considered partner traits have found that imperfect sex discrimination in males is not random, but rather varies depending on individual-level variation in the traits of conspecifics (Han et al., 2016; Mizumoto et al., 2022; Richardson and Zuk, 2023). The question of which traits are used to identify sexes connects to larger questions of how individuals choose mating partners—a central issue in selection selection theory (DuVal et al., 2023; Rosenthal, 2017). Interestingly, we find that in some cases selection may favor degradation of signal quality, as a result of selective pressure for a lower mating rate. This result relates to the large body of work on sexual conflict over the mating rate (Parker, 2006). Here, the low-investment sex is generally expected to favor a higher mating rate because additional matings lead to additional fertilizations, whereas the high-investment sex is expected to favor a lower mating rate because additional matings incur costs without increasing fecundity. In our model, when mating opportunities are both plentiful and costly, Signalers may experience selection for lower signal quality (i.e., obfuscated sexual identity) as a mechanism to avoid overmating. Lower signal quality, in turn, removes the benefit of sex discrimination and thus leads to its loss and an increase in the expression of SSB. That is, Signaler evolution drives selection for Finder ISB. This trajectory is reminiscent of biological scenarios in which females plastically display male-like traits to reduce their mating rate (Willink et al., 2019). For example, in *Littorina* snails, females that live in highdensity conditions stop providing a sex-specific cue in their mucus, leading to males being unable to preferentially follow female mucus trails, as they do in low-density conditions (Johannesson et al., 2010).

### Effects of encounter rate

Our model is the first to explicitly consider the role of the encounter rate in shaping the evolution of sex discrimination. Empirical studies have assessed how different encounter rates lead to differences in the expression of SSB, with conflicting results. For example, SSB in the deep-sea squid (*Octopoteuthis deletron*) has been used to argue that low encounter rates lead to high costs of missed matings which, in turn, lead to low sex discrimination (Hoving et al., 2012). Similarly, social isolation has been shown to increase the rate at which males engage in SSB in fruit flies (*Drosophila melanogaster*) (Bailey et al., 2013; Dukas, 2010), burying beetles (*Nicrophorus vespilloides*) (Engel et al., 2015), and red flour beetles (*Tribolium castaneum*) (Martin et al., 2015). Outside of SSB-focused studies, non-reproductive mating has also been shown to increase in response to low encounter rates in a model of female multiple mating (Kokko and Mappes, 2012). However, this result appears not to be universal, as some evidence exists that high density conditions (which presumably correspond to increased encounter rates) lead to increased expression of SSB (He, 2008; Livingstone and Ramani, 1978; Weldon, 2005). High encounter rates may lead to weak sex discrimination if males engaged in intense scramble competition forego sex discrimination to mate as quickly as possible (Han and Brooks, 2015). In sum, there is some evidence that both very low (social isolation) and very high (high density) encounter rates lead to weak sex discrimination (Scharf and Martin, 2013).

Likewise, we also find that both very low and very high encounter rates lead to more SSB (i.e., larger basin of attraction for the ISB equilibrium). For low encounter rates, our results directly support the hypothesis that high opportunity costs to missed matings given low encounter rates lead to weak sex discrimination (e.g., in deep-sea squid; Hoving et al., 2012). For higher encounter rates, while our model appears to capture empirically observed patterns, there are key differences. Many of these past empirical studies were assessing not the evolution of sex discrimination, but plastic responses to different social conditions such as density and sex ratio. Further, signals of sexual identity were not considered in these empirical studies, but are critical to our result that high encounter rates increasingly favor ISB. Under such high encounter rates, high-investment Signalers are readily mated (i.e. there is no limitation of Finder gametes) and therefore reduce signaling to avoid excess costly mating. This appears to be a different mechanism from the SSB arising from scramble competition in male-biased experimental populations. Further differences emerge on the role of encounter rate in the investment-role-reversed case.

Here, the DSB basin of attraction expands monotonically as the encounter rate increases. The difference in the two investment-role cases arises from an asymmetry between Finders and Signalers in our model. In both cases, when encounters are frequent, the high-investment sex experiences selection for a lower overall number of matings. However, Signalers and Finders evolve lower mating rates through different mechanisms. Signalers experience selection for a lower quality signal, making SSB more likely. Finders, on the other hand, experience selection for increased sex discrimination, making SSB less likely. Consistent with the investment-role-reversed case, past work on opposite-sex mate choice has shown increasing choosiness with increasing encounter rates (Gowaty and Hubbell, 2009).

### Mating costs

Our model incorporates explicit costs of mating, in contrast to previous models that assign costs directly to sex discrimination (Lerch and Servedio, 2021, 2023) or assume a tradeoff between mate targeting and gamete production (Parker, 2014). These costs may reflect metabolic expenditure, time, risk of injury, or other trade-offs arising from the act of mating. Interestingly, the consequences of mating costs differ depending on which sex they arise in. Increasing costs for Finders results in more discrimination (leading to less SSB), while increasing costs for Signalers results in signal degradation (leading to more SSB). These findings highlight mating costs as a key variable influencing the evolution of SSB.

Although direct costs to mating are often assumed in theoretical arguments, the prevalence of non-reproductive copulatory behavior—same-sex, different-sex, masturbatory (Sommer et al., 2022), necrophilic (Toyoda et al., 2024), or cross-species (Depret et al., 2025)—suggest that, for many animals, such costs may be marginal when incorporated into a calculation of total lifetime fitness. Most of the empirical literature on costs of mating refers to factors that are either beyond the scope of our model (e.g., post-birth parental care) or accounted for elsewhere in our model; e.g., gametic investment costs reflected in our investment roles, and sexual ornamentation costs captured in the signaling cost *C*(*q*). Studies that do evaluate the direct costs of mating vary in their findings. Some studies have documented survival costs of mating through either decreases in longevity (Fedorka et al., 2004; Martin and Hosken, 2004; Pereira et al., 2010; Stojkovic et al., 2010) or increased likelihood of predation (Magnhagen, 1991). Other studies have found costs of signaling but not mating (Cordts and Partridge, 1996; Papadopoulos et al., 2010). Still other studies have found no longevity costs at all (Ferkau and Fischer, 2006; Perez-Staples and Aluja, 2006), or that overall fitness benefits of multiple mating outweigh any survival costs, possibly even beyond an increase in fertilization rates (Rodrigues et al., 2020; Worthington and Kelly, 2016). In sum, the prevalence and significance of mating costs appear to be highly species- and context-specific. In particular, the negligible-cost case—with a consequently large basin of attraction for ISB—is likely relevant to many biological systems.

### Sex ratio

In general, we find that Finder-biased sex ratios favor the evolution of sex discrimination. In populations with Signaler-biased sex ratios, Finders can mate indiscriminately and most matings will be with Signalers by chance. This result matches past theory on the evolution of sex discrimination (Lerch and Servedio, 2021) and also a study of experimental evolution in red flour beetles (*Tribolium castaneum*) (Sales et al., 2018). Interestingly, as noted by Lerch and Servedio (2023), many studies of plastic sex discrimination find a different pattern, such that Finder-biased sex ratios do not lead to stronger sex discrimination (Engel et al., 2015; Han and Brooks, 2015; Macchiano et al., 2018; Svetec and Ferveur, 2005; Switzer et al., 2004). While outside the scope of our study, this discrepancy suggests that theoretical modeling of fixed versus plastic sex discrimination would be a valuable contribution.

### Ancestral conditions and SSB

As was first pointed out by Parker (2014), there is no reason to think that sex discrimination was an immediate byproduct of the evolution of two sexes. Rather, indiscriminate mating was most likely the ancestral mode of sexual behavior for early animals (Monk et al., 2019). The existence of a stable ISB equilibrium suggests that ancestral ISB need not have been immediately followed by the evolution of strong sex discrimination. Instead, without detectable differences between sexes (which might arise, for example, as a byproduct of anisogamy) ancestral ISB could persist for indefinite stretches of evolutionary time.

This does not necessarily imply, however, that contemporary SSB is always derived from ancestral ISB (Dickins and Rahman, 2020). Sexual behaviors of early animals (likely broadcast spawners) are different from derived sexual behaviors (e.g., courtship behaviors in insects). In many cases, SSB does not appear to be indiscriminate, and may evolve for other reasons (Kureck et al., 2011; Lane et al., 2016; Levan et al., 2009; Mizumoto et al., 2022; Preston-Mafham, 2006). Moreover, contemporary instances of apparent ISB (Hoving et al., 2012; Marco and Lizana, 2002; McCarthy and Young, 2002; Slattery and Bosch, 1993; Young et al., 1992) cannot be clearly traced to a trait in a common ancestor given their diverse forms of expression. Our model shows that both ancestral and contemporary ISB can persist under natural selection, providing a plausible explanation for many occurrences of SSB.

The question then arises as to how ancestral ISB could transition to DSB, or vice versa. Such transitions might occur via stochastic drift, environmental change, or byproducts of other traits. For example, at ISB equilibrium, a signal could arise stochastically or as a byproduct of repoductive physiology, bringing the population into the DSB basin of attraction. Conversely, shifts in prevailing environmental conditions could lead to a degradation in signal quality, which, in turn, could result in transition from DSB to ISB equilibrium. Similar dynamics often play out in the loss of extrinsic barriers to reproductive isolation in cases of “reverse speciation” (Frei et al., 2022; Seehausen, 2006; Seehausen et al., 1997; Zhang et al., 2019).

For spatially subdivided populations, our results suggest that differences in mating ecology parameters across subpopulations could lead to spatial heterogeneity in observed sexual behavior. Different populations could evolve towards different equilibria (ISB vs. DSB) based on differences in conditions such as density or sex ratio. Moreover, high levels of dispersal or migration could lead to the maintenance of alternative mating strategies within populations.

## Conclusions

Our results demonstrate that the coevolution of sex discrimination and sexual signals often produces bistable dynamics, with both exclusive DSB and exclusive ISB as stable selective equilibria. Given that our modeling framework and assumptions are significantly different from a previous study also finding bistability of signaling and discrimination (Lerch and Servedio, 2023), this result appears robust to system-specific details. We find that low encounter rates favor ISB because of the high opportunity cost of missing a mating. Higher encounter rates may lead to either ISB or DSB, depending on investment roles, mating costs, the nature of discrimination and signaling. Broadly, our modeling approach reveals the role of key ecological parameters in determining whether natural selection leads to discriminate or indiscriminate sexual behavior.

Our model adds to growing theoretical support for the hypothesis that some instances of SSB are indiscriminate in nature. In particular, this hypothesis applies to courtship and copulatory behaviors in taxa for which incomplete distinguishability between sexes is plausible, either as an ancestral or derived state. Other instances of SSB, such as pair-bonding or co-parenting in birds or mammals, likely involve other explanations. Indeed, as SSB is an umbrella term encompassing a wide variety of behaviors, it is likely that a variety of evolutionary forces have shaped these behaviors. Moreover, as Parker (2014) and Monk et al. (2019) make explicit, exclusively different-sex behavior—that is, avoidance of SSB—is itself an evolved behavior requiring explanation where it arises. Ultimately, understanding the rich diversity of mating and reproductive behavior demands careful observation and modeling of the relevant ecological and evolutionary dynamics.

## Acknowledgments

We are grateful to Ambika Kamath, Anuraag Mukherjee, Maria Servedio, Delaney O’Connell, Nancy Emery, Alec Chiono, Katherine Bardsley, and Miles Moore for conversations and feedback. B.A. was supported by grants #62220 and #63454 from the John Templeton Foundation. B.A.L. was supported by a Royster Fellowship from the UNC Graduate School.

## A Derivation for arbitrary sex ratio

Here we derive the invasion fitness functions and adaptive dynamics equations for our model in the case of arbitrary sex ratio, as quantified by the fraction *s* of Signalers. This generalizes the case of even sex ratio (*s* = 1/2) derived in the main text. These invasion fitness functions quantify the growth rate of a rare mutant type of discrimination strength *d*′ (for Finders) or signal quality *q*′ (for Signalers) in a resident population with trait values *d* and *q*.

We break into cases according to which sex is high-investment. The high-investment sex is fertilized the first time they mate with the opposite sex in a given season, and cannot be fertilized again in the same season (although they remain available for nonreproductive mating). The low-investment sex fertilizes as many opposite-sex conspecifics as they are first to mate with in a given season.

### A.1 High-investment Signalers (baseline case)

We first suppose that Signalers are high-investment. To derive invasion fitness for Signalers, consider a mutant Signaler type, with signal quality *q*′. This Signaler encounters Finders at rate 2(1 − *s*)*λ*, and each encounter results in mating with probability 1 − (1 − *q*′)*d*, according to Eq. (1). So overall, this Signaler experiences mating as a Poisson process with rate 2(1 − *s*)*λ*(1 − (1 − *q*′)*d*), obtained by combining Eq. (1) with the Poisson process encounter model. The probability that this Signaler is fertilized (mates at least once) in a given season is 1 − *e*^−2(1 − *s*)*λ*(1 − (1 − *q′*)*d*)^, yielding expected benefit *b* 1 − *e*^−2(1 − *s*)*λ*(1 − (1 − *q′*)*d*)^. Combining this with the costs of mating and signaling, the invasion fitness of the mutant Signaler type is

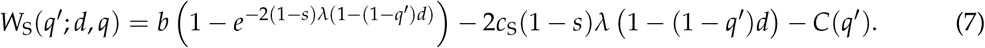

Now consider a mutant Finder of discrimination strength *d*′. A given resident Signaler experiences mating from this Finder at rate (*λ*/*N*)(1 − (1 − *q*)*d*′), and from resident Finders at total rate 2(1 − *s*)*λ*(1 − (1 − *q*)*d*). The probability that the mutant Finder is the first to mate with this Signaler is

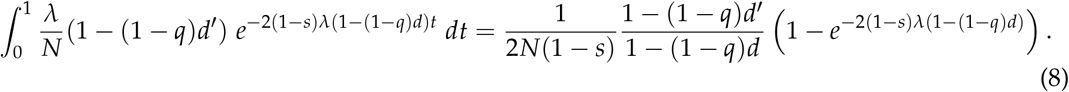

Multiplying by the number of Signalers, 2*Ns*, the expected number of fertilizations per time-step achieved by this Finder is

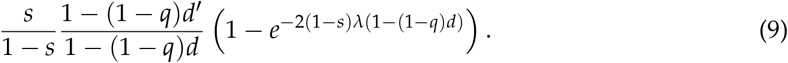

Meanwhile, this mutant Finder mates with Signalers at rate 2*λs*(1 − (1 − *q*)*d*′), and with Finders at rate 2*λ*(1 − *s*)*λ*(1 − *d*′), incurring a total cost of 2*c*_F_*λ*(1 − (1 − *sq*)*d*′). Aggregating benefits and costs, we obtain the invasion fitness of the mutant Finder type:

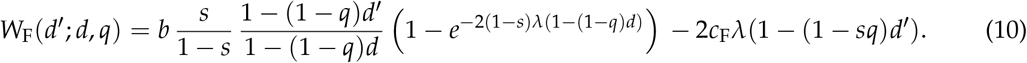

Applying the adaptive dynamics approach, we suppose that mutations in signal quality and discrimination strength occur at respective rates *µ*_S_ and *µ*_F_ (assumed small), with expected step size *σ*_S_ and *σ*_F_ (also assumed small). According to the canonical equation of adaptive dynamics, long-term trait dynamics are characterized by

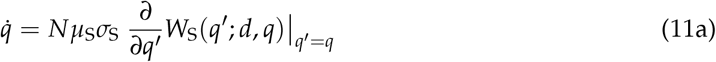

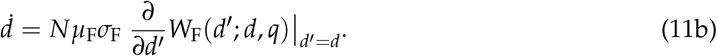

Taking the appropriate derivatives of Eqs. (7) and (10), we obtain

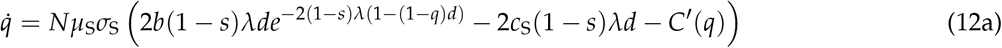

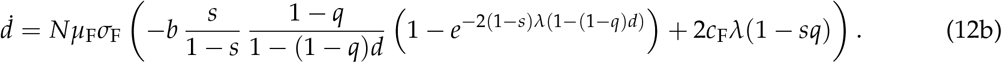

For even sex ratio (*s* = 1/2), we recover Eq. (5) of the main text.

### A.2 High-investment Finders (role-reversed case)

We now reverse roles, so that Finders are high-investment. To compute invasion fitness for Finders, consider a mutant Finder of discrimination strength *d*′. This Finder mates with Signalers at rate 2*sλ*(1 − (1 − *q*)*d*′). The probability of at least one mating per season is therefore 1 − *e*^−2*sλ*(1 − (1 − *q*)*d′*^). This Finder also mates with other Finders at rate, 2(1 − *s*)*λ*(1 − *d*′), yielding a total mating cost of 2*c*_F_*λ*(1 − (1 − *sq*)*d*′). Aggregating benefits and costs, we obtain the invasion fitness

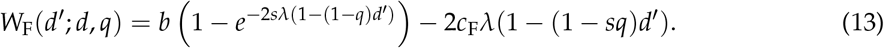

Now, consider a mutant Signaler, with signal quality *q*′, and a given resident Finder. This Finder mates with this mutant Signaler at rate (*λ*/*N*) (1 − (1 − *q*′)*d*), and with resident Signalers at total rate (2*sλ*) (1 − (1 − *q*)*d*). The probability that the mutant Signaler is the first to mate with this Finder is

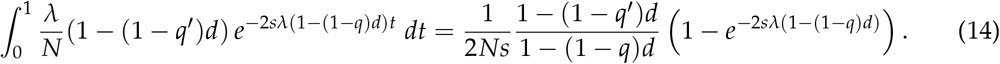

Multiplying by the total number of Finders, 2*N*(1 − *s*), the expected number of fertilizations for this mutant Signaler is

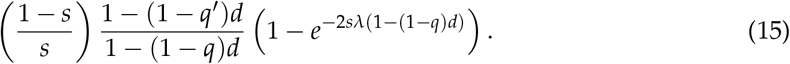

Since this Signaler mates with Finders at rate 2(1 − *s*)*λ*(1 − (1 − *q*′)*d*), the incurred cost is 2*c*_S_(1 − *s*)*λ*(1 − (1 − *q*′)*d*). Combining cost and benefits, we obtain the invasion fitness

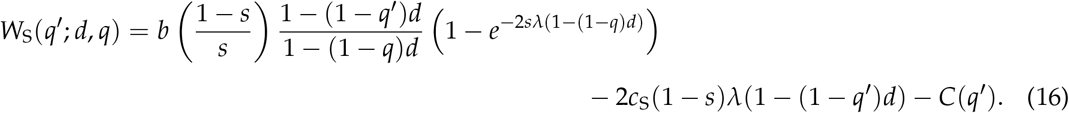

The adaptive dynamics equations become

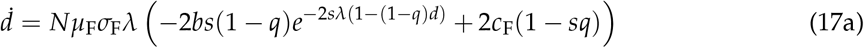

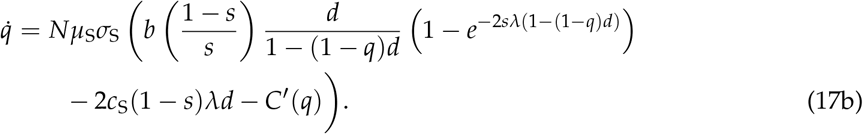

Substituting *s* = 1/2, we recover Eq. (6) of the main text.

## B Stability of equilibria

Here we obtain conditions for the stability of equilibria. We confine our analysis to the baseline investment case; the role-reversed case can be analyzed similarly.

We consider two notions of stability. A fixed point 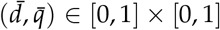 is *convergence stable* if it is a stable fixed point of the adaptive dynamics, Eq. (12). Fixed point 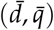 is *evolutionarily stable* if it is robust to invasion by mutant types, meaning 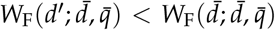 and 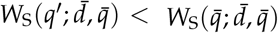 for all *d*′ and *q*′. Neither of these conditions necessarily implies the other; see Geritz et al. (1998) for theoretical background.

Our results on stability can be summarized in the following theorem:

### Theorem 1.

*In the baseline investment case, suppose that c*_F_ > 0 *and C*(*q*) *is linear with slope C*′ > 0.

*Then*

i. *The pure ISB state of p* = *q* = 0 *is both convergence stable and evolutionarily stable if bs* 1 − *e*^− 2(1 − *s*)*λ*^ > 2(1 − *s*)*λc_F_*;
ii. *The pure DSB state of p* = *q* = 1 *is both convergence stable and evolutionarily stable if* 2*λ*(1 − *s*) *be*^− 2(1 − *s*)*λ*^ − *c*^S^ > *C*′;
iii. *If (i) and (ii) both hold, the adaptive dynamics have a single interior fixed point, which is an unstable saddle point*.

We prove claims (i)-(iii) in the subsections below. It is not always necessary to assume *C*(*q*) is linear; this assumption is relaxed in various ways for specific results. Substituting *s* = 1/2 for even sex ratio, the stability conditions presented in the main text are recovered.

Our proofs make use of the following function:

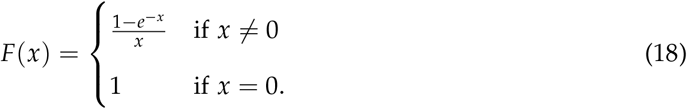

With *F*(*x*) defined as above, the condition in claim (i) of Theorem 1 can be rewritten as *bsF* 2(1 − *s*) *λ*) > *c*_F_.

We also define the quantity *r* = 2(1 − *s*)*λ*(1 − (1 − *q*)*d*), which represents the rate at which a given Signaler experiences mating attempts from Finders. Setting *Nµ*_S_*σ*_S_ = *Nµ*_F_*σ*_F_ = 1 without loss of generality (since these do not affect locations or stability of equilibria), Eq. (12) for adaptive dynamics in the baseline case can then be rewritten as

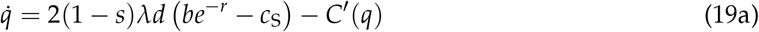

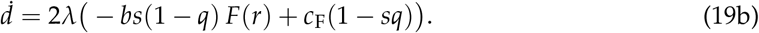

The quantity *F*(*r*) appearing in Eq. (19b) quantifies the efficacy of Finder mating attempts; that is, the probability 1 − *e*^− *r*^ that a given attempt with a Sender results in fertilization, normalized by the rate *r* at which such attempts are made.

We establish the following properties of *F*(*x*):

### Lemma 1.

*F*(*x*), *as defined in Eq*. (18), *is continuous, differentiable, and decreasing for all real values of x*.

*Proof*. Continuity and differentiability follow from the Taylor series

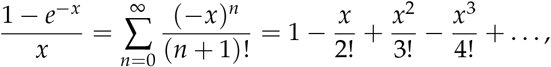

which converges at all real *x*.

The derivative of *F*(*x*) is

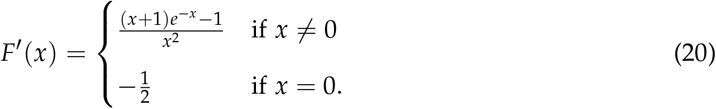

The derivative of the numerator in the *x* ≠ 0 case of Eq. (20) is − *xe*^− *x*^, which is positive for *x* > 0 and negative for *x* <0. It follows that the numerator attains a maximum of zero at *x* = 0 and is otherwise negative. Since the denominator is positive for all *x* ≠ 0, it follows that *F*′(*x*) <0 for all *x*, i.e. *F*(*x*) is decreasing everywhere.

### B.1 ISB equilibrium

We first establish claim (i) of Theorem 1, regarding the pure ISB state of no signaling or discrimination (*d* = *q* = 0). This state is a convergence stable equilibrium if 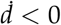 and 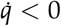 at this point. From Eq. (19), this occurs if *bsF (*2(1 − *s*)*λ)* > *c*_F_ and *C*′(0) > 0. The first condition says that a Finder’s expected benefit of mating exceeds its cost, given that, under indiscriminate mating, a fraction *s* of a Finder’s matings will be with other Finders. The second condition says that signaling is costly, even when of low quality.

These two conditions also suffice to guarantee evolutionary stability. To see this, we evaluate Eqs. (7) and (10) to obtain

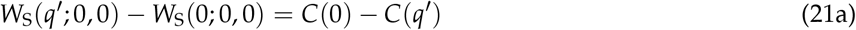

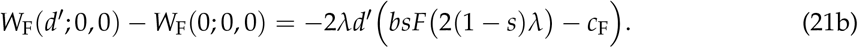

The condition *C*′(0) > 0—together with our assumption that *C*(*q*) is nondecreasing—guarantees that the right-hand side of Eq. (21a) is negative for all *q*′ > 0. Likewise, the condition *bsF* (2(1 − *s*)*λ* > *c*_F_ guarantees that the right-hand side of Eq. (21b) is negative for all *d*′ > 0. Thus, if these two conditions are met, the ISB equilibrium is uninvadable by any mutant types. This proves claim (i).

### B.2 DSB equilibrium

We now establish claim (ii). The pure DSB state of perfect signaling and discrimination (*d* = *q* = 1) is a convergence stable equilibrium if 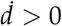 and 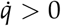 at this point. From Eq, (19), this occurs if and only if *c*_*F*_ > 0 and 2*λ*(1 − *s*) (*be*^− 2(1 − *s*)*λ*^ − *c*_S_) > *C*′(1). The second condition asserts that the net benefits gained from mating exceed the cost of maintaining perfect signal quality.

These two conditions, together with an additional assumption that *C*(*q*) is linear or convex (i.e. the marginal cost of signal quality is constant or increasing), suffice to guarantee evolutionary stability as well. We prove this as a separate theorem:

#### Theorem 2.

*Under the conditions that*

i. *c*_F_ > 0,
ii. 2*λ*(1 − *s*) *be*^− 2(1 − *s*)*λ*^ − *c*_S_ > *C*′(1), *and*
iii. *C*(*q*) *is linear or convex, the DSB state d* = *q* = 1 *is evolutionarily stable*.

*Proof*. We begin by computing

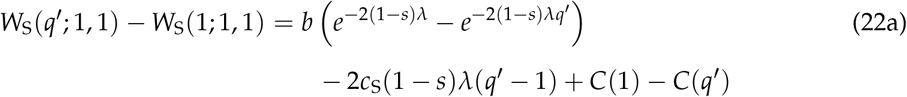

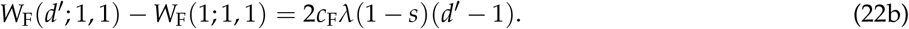

From Eq. (22b), we observe that Condition (i) implies *W*_F_(*d*′; 1, 1) <*W*_F_(1; 1, 1) for all *d*′ <1. Turning to Eq. (22a), we rewrite this equation as

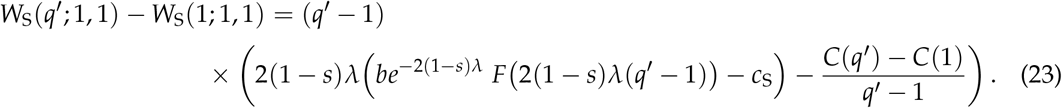

Since *F*(*x*) is decreasing in *x*, we have *F (*2(1 − *s*)*λ*(*q*′ − 1)) > 1 for all *q*′ <1. Meanwhile, under Condition (iii), we have (*C*(*q*′) − *C*(1)) /(*q*′ − 1) ≤ *C*′(1) for *q*′ <1. Combining these with Condition (ii), we have

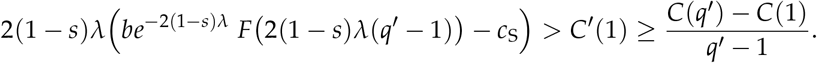

The right-hand side of Eq. (23) is therefore negative, and hence *W*_S_(*q*′; 1, 1) <*W*_S_(1; 1, 1), for all *q*′ <1.

This establishes claim (ii) of Theorem 1.

### B.3 Unstable interior fixed point

We now move to claim (iii), which we also prove as a separate theorem:

#### Theorem 3.

*Under the conditions that*

i. *c*_F_ > 0,
ii. *bs* 1 − *e*^− 2(1 − *s*)*λ*^ > 2(1 − *s*)*λc*_F_,
iii. 2*λ*(1 − *s*) *be*^− 2(1 − *s*)*λ*^ − *c*_S_ > *C*′(1), *and*
iv. *C*(*q*) *is linear or concave*,

*the adaptive dynamics, Eq*. (19), *have a single interior saddle point*.

*Proof*. We first show that, if an interior fixed point exists, it must be an unstable saddle point. For this, we compute the Jacobian matrix:

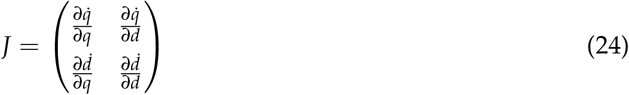

The relevant partial derivatives are:

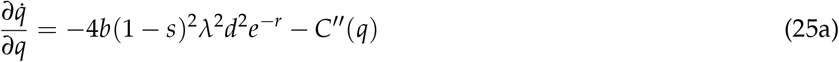

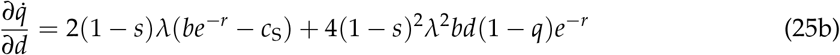

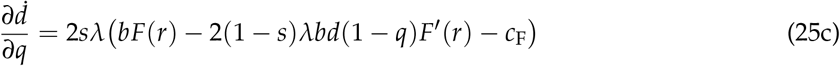

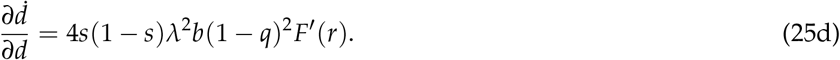

The determinant, det 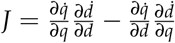, simplifies to

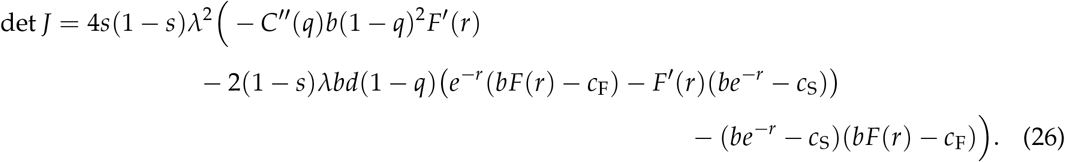

We will show that det *J* <0 everywhere. We first observe that Conditions (ii) and (iii) imply *bF*(2(1 − *s*)*λ*) > *c*_F_ and *be*^− 2(1 − *s*)*λ*^ > *c*_F_. Since *r* attains a maximum of 2(1 − *s*)*λ* at *q* = 1, and *e*^− *x*^ and *F*(*x*) are decreasing functions, it follows that *bF*(*r*) > *c*_F_ and *be*^− *r*^ > *c*_S_ for all (*d, q*) ∈ [0, 1] × [0, 1]. Meanwhile, *C*^*′′*^(*q*) ≤ 0 by Condition (iv). Applying these observations to Eq. (26), we obtain that det *J* <0 for all (*d, q*) ∈ [0, 1] × [0, 1]. This means that *J* always has one positive and one negative eigenvalue. We conclude that any interior fixed point of the adaptive dynamics must be an unstable saddle point.

To prove that exactly one such interior fixed point exists, we make use of the Lefshetz Fixed Point Theorem (see section 3.4 of Guillemin and Pollack, 1974). Let *ϕ*_*t*_ : [0, 1] × [0, 1] → [0, 1] × [0, 1] denote the flow of the adaptive dynamics equations (projecting onto boundaries). *ϕ*_0_ is the identity mapping, and is homotopic to *ϕ*_*t*_ for any *t* > 0. It then follows from the Lefshetz Fixed Point Formula that the sum of the indices of all fixed points equals the Euler characteristic of the space, which for the square [0, 1] × [0, 1] is 1. Stable fixed points have an index of 1, while saddle points have an index of −1. Since all interior fixed points must be saddle points, if there are two stable fixed points, there must be exactly one interior saddle point.

This completes the proof of Theorem 1.

## C Small encounter rates

An extreme case of our model is the rare encounter regime, *λ* ≪ 1. In this regime, the opportunity cost of missed mating is very high, since the likelihood of encountering multiple potential mates in a season is negligible.

In this regime, our model simplifies to a linear system that can be solved exactly. Using the approximation *e*^− *x*^ ≈ 1 − *x*, the adaptive dynamics equations for our model reduce to

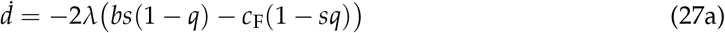

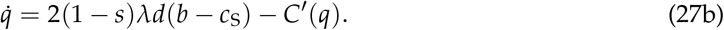

This result applies to both cases of investment roles; i.e. the investment-role cases become equivalent in the rare encounter regime. This is because investment roles come into play only when there is a significant likelihood of multiple opposite-sex matings per season.

**Supplementary Figure S1:**
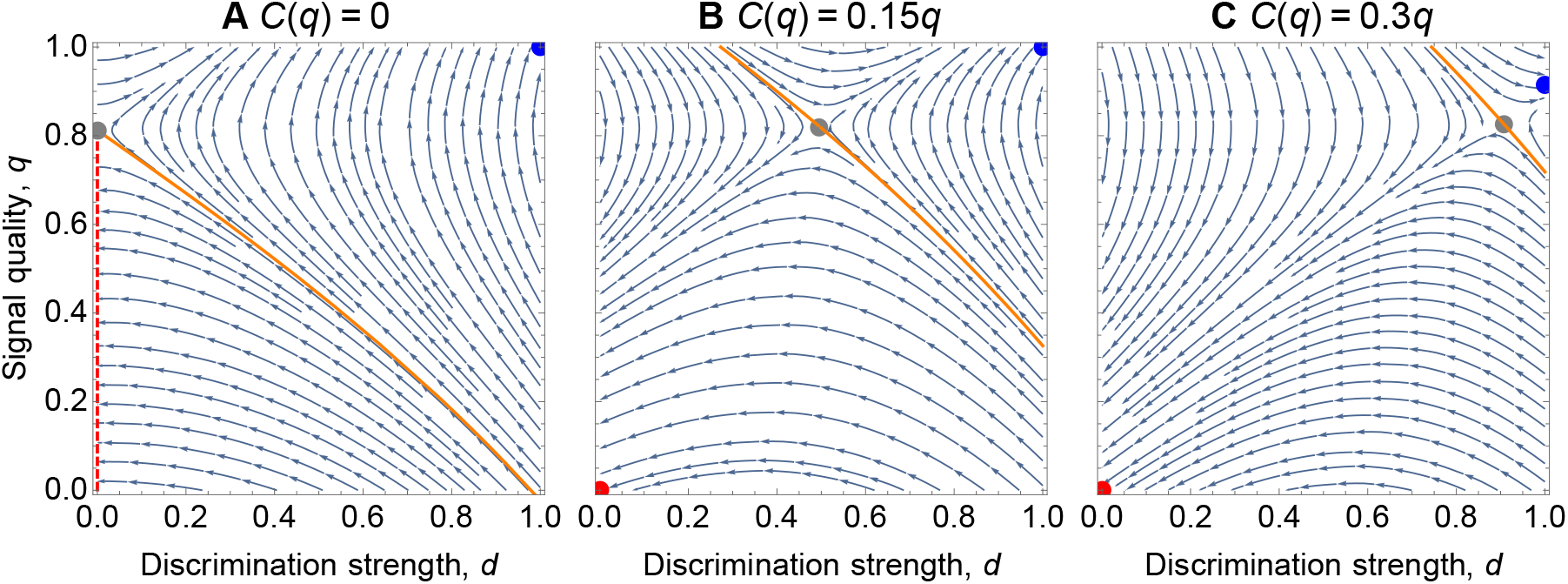
Effect of signaling costs. We varied the signaling cost *C*(*q*) as a function of signal quality, *q*, with other parameters fixed at *b* = *λ* = 1, *c*_F_ = *c*_S_ = 0.1. **A** If signaling is costless (or arises as a byproduct of other traits), indiscriminate sexual behavior (*d* = 0) can still evolve, but at a range of neutrally stable signal qualities (red dashed line) instead of a single ISB equilibrium. This is because, in the absence of discrimination and signal costs (*d* = *C*(*q*) = 0), there is no selection on signal quality 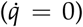. **B** For positive signaling costs, a stable ISB equilibrium appears. **C** As its costs increase, signaling becomes less advantageous, and consequently, the ISB basin of attraction expands.

An interior fixed point must satisfy 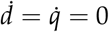. Solving, we obtain

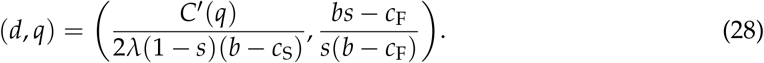

If both coordinates above are between 0 and 1, then the interior fixed point exists—and is an unstable saddle point according to Theorem 3—and the pure ISB and DSB points are both stable fixed points. Note that this requires *C*′(*q*) to be of comparable magnitude to *λ* (i.e. the marginal cost of signaling must be comparably small to the encounter rate), as well as *b* > *c*_S_ and *bs* > *c*_F_.

At the interior fixed point (if it exists), the eigenvalues of the Jacobian are 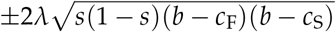.

The corresponding eigenvectors are

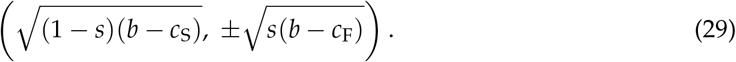

The separatrix between the basins of attraction of the ISB and DSB equilibria is the line with direction vector 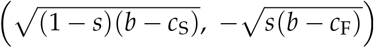 (the eigenvector corresponding to the negative eigenvalue) passing through the interior fixed point.

**Supplementary Figure S2:**
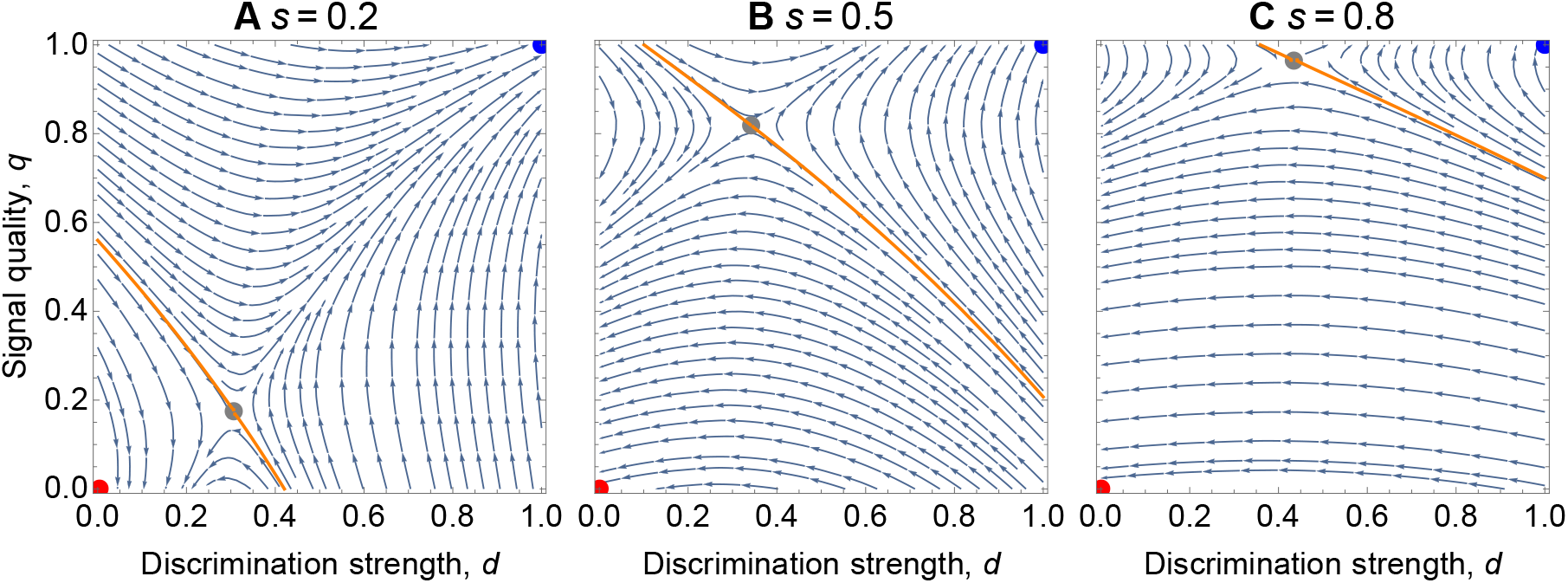
Effect of sex ratio. We varied the fraction, *s*, of Signalers in the population, with other parameters fixed at *b* = *λ* = 1, *c*_F_ = *c*_S_ = *C*′(*q*) = 0.1. **A** Under a Finder-biased sex ratio (*s* <0.5), Finders mostly encounter each other. Selection favors discrimination to avoid nonreproductive Finder-Finder matings, leading trajectories to DSB equilibrium. **B** As Signalers become more abundant, discrimination becomes less necessary. **C** For a Signaler-biased sex ratio (*s* > 0.5), Finders mostly encounter the opposite sex. There is little advantage to be gained from discrimination, so most trajectories lead to ISB.

## Notes

### Competing Interest Statement

The authors have declared no competing interest.

